# Fast activation cycles of Rac1 at the lamellipodium tip trigger membrane protrusion

**DOI:** 10.1101/130849

**Authors:** Amine Mehidi, Olivier Rossier, Anaël Chazeau, Fabien Binamé, Amanda Remorino, Mathieu Coppey, Zeynep Karatas, Jean-Baptiste Sibarita, Violaine Moreau, Grégory Giannone

## Abstract

The spatiotemporal coordination of actin regulators in the lamellipodium determines the dynamics and architecture of branched F-actin networks during cell migration. The WAVE complex, effector of Rac1 during cell protrusion, is concentrated at the lamellipodium tip. Yet, correlation of Rho GTPases activation with cycles of membrane protrusions, suggested that Rac1 activation is not synchronized with membrane protrusion and occurs behind the lamellipodium. However, RhoA activation is maximal at the cell edge and synchronized with edge progression. Combining single protein tracking (SPT) and super-resolution imaging with loss- or gain-of-function of Rho GTPases mutants, we demonstrate that Rac1 immobilizations at the lamellipodium tip are correlated with Rac1 activation, on the contrary to RhoA. We show that Rac1 effector WAVE and Rac1 regulator IRSp53 accumulate at the lamellipodium tip by membrane free-diffusion and trapping. Nevertheless, wild-type Rac1, which directly interacts with WAVE and IRSp53, only displays slower diffusion at the lamellipodium tip, suggesting fast local activation/inactivation cycles. Local optogenetic activation of Rac1, triggered by Tiam1 membrane recruitment, proves that Rac1 activation must occur at the lamellipodium tip and not behind the lamellipodium to trigger efficient membrane protrusion. Furthermore, coupling tracking with optogenetic activation of Rac1 demonstrates that Rac1-WT diffusive properties are unchanged despite enhanced lamellipodium protrusion. Taken together, our results support a model where Rac1 is rapidly switching between activation and inhibition at the lamellipodium tip, ensuring a local and fast control of Rac1 actions on its targets.

**Significance:** Rac1 and RhoA GTPases are molecular switches controlling the actin cytoskeletal during cell migration. WAVE, Rac1 effector during cell protrusion, is concentrated at the lamellipodium tip. But, recent biosensor imaging studies suggested that Rac1 activation occurs behind the lamellipodium, while RhoA activation is maximal at the cell edge. Using single-molecule imaging and optogentics Rac1 activation we solved this apparent contradiction. We revealed a strong correlation between Rac1 activation and transient immobilizations at the lamellipodium tip, unlike RhoA. Furthermore, we demonstrated that Rac1 must be activated at the lamellipodium tip and not away from it to stimulate protrusion. Thus, fast cycling between activation and inhibition at the proximity of Rac1 targets ensures a local and fast control over Rac1 actions.

**Abbreviations:**

Arp2/3: actin related proteins 2/3
D: diffusion coefficient
F-actin: actin filaments
FMNL2: formin-like protein-2
FN: fibronectin
GAP: GTPase-activating protein
GDI: Guanine-nucleotide Dissociation Inhibitor
GEF: Guanine-nucleotide Exchange Factor
IRSp53: insulin receptor tyrosine kinase substrate p53
LM: lamellipodium
NPF: nucleation promoting factor
MSD: mean squared displacement
PALM: photoactivation localization microscopy
PSD: post synaptic density
rconf: confinement radius
spt: single protein tracking
VASP: vasodilator-stimulated phosphoprotein
WAVE: WASP-family verprolin homologue

## Introduction

Cell motility is an integrated process involved in critical physiological phenomenon such as embryogenesis, immunological response, wound healing, and growth cone path finding (1). Dysregulation of cell motility, which has a precise hierarchical spatial and temporal organization, contributes to pathologies including tumor formation and metastasis (2). At the cellular (3), sub-cellular (4, 5), and molecular (1, 6) levels, migration proceeds through cycles lasting from minutes to seconds. During those cycles, macromolecules undergo motions and transient interactions that are essential to their function, and this is particularly true during cell migration where nanomachine-like protein complexes involved in adhesion and actin polymerization must be at a specific location at a precise time (7, 8).

The first step in cell motility is the forward protrusion of the lamellipodium which is a thin sheet of membrane-enclosed F-actin networks propelled by actin polymerization (7, 9). Rho GTPases are acting as molecular switches to control cytoskeletal rearrangements during cell growth, adhesion and motility (10). Among them, Rac1 activation is the major signal driving lamellipodium formation (11–13). Rac1 is targeted to the plasma membrane where it can bind and activate the WAVE complex, a nucleation promoting factor (NPF) that stimulates the Arp2/3 complex and thus nucleation of branched F-actin (14, 15). F-actin length is then controlled by elongation factors including VASP and Formin-Like protein (FMNL2) (16, 17) and capping proteins (18). Finally, F-actin turnover is driven by severing proteins such as ADF/cofilin (19, 20) or myosin motors (21, 22). The actions of actin regulators in the lamellipodium are highly compartmentalized (20, 23). Actin nucleation, branching and elongation occur at the lamellipodium tip membrane (9), myosin II generates forces at the back of the lamellipodium (5), ADF/cofilin ensures F-actin severing within the entire lamellipodium (20). From the complex spatiotemporal coordination of actin regulators at the nanometer scale could emerge the micron-scale periodic phenomenon driving lamellipodium protrusion (4, 24).

The development of super-resolution microscopy techniques (25, 26) and single protein tracking (SPT) (8, 27) has revolutionized biomolecular imaging in cells. The direct observation of biomolecules at the single molecule level has enabled their localization and tracking at the nanometer scale (28, 29). Super-resolution microscopies based on single molecule localization have been used to study sub-cellular structures including, integrin-based adhesion sites (6, 30, 31), neuronal axons (32) and dendritic spines (33, 34). Those studies unraveled that proteins, which seem co-localized using conventional light microscopy, are spatially segregated at the nanoscale level into distinct functional layers. The lamellipodium is also a polarized structure organized in functional domains, the tip, the core and the back. The localization of most actin binding proteins and regulators in the lamellipodium, obtained using conventional fluorescent microscopy, are in good agreement with their functions (5, 20, 23, 35). Nevertheless, the sites of action of signaling proteins are often difficult to pinpoint. For more than a decade, Rac1 was associated with lamellipodium formation (11–13). Since WAVE is concentrated at the lamellipodium tip (20, 36), activated Rac1 should work at this location to stimulate WAVE. However, FRET-based experiments, which correlated Rac1 and RhoA activities with cycles of membrane protrusions, suggested that Rac1 is activated 2 μm away from the lamellipodium tip (37). Thus, Rac1 activation seemed not synchronized with membrane protrusion on the contrary to RhoA activation which is located at the cell edge (38) and synchronized with edge progression (37). Thus, the spatial resolution of conventional fluorescent microscopy appears insufficient to faithfully decrypt the sequence of molecular events controlling actin network dynamics within protrusive structures. In this study, we used SPT and super-resolution microscopy, which give us molecular resolution on the localization of Rac1 and RhoA inside and outside the lamellipodium. Combining these techniques with loss- or gain-of-function of Rho GTPases mutants, we linked the nanoscale organization and dynamics of Rac1 to its activation and site of action within the lamellipodium. In addition, we used optogenetics to trigger spatially controlled activation of Rac1 and study the consequences on Rac1 molecular dynamics and membrane protrusion.

## Results

### The WAVE complex and IRSp53 are recruited at the lamellipodium tip by membrane free-diffusion

Nucleation of F-actin branches requires the coordination in space and time of different signals involving GTP-bound Rac1, the WAVE and Arp2/3 complexes, IRSp53 and acidic phospholipids (PIP3) (14, 15, 39). However, understanding their coordination within the lamellipodium implies knowledge of their molecular dynamics. To determine the sequence of molecular events leading to Rac1-dependent WAVE activation, we performed high-frequency sptPALM acquisition (50 Hz) to characterize the diffusive properties of actin regulators in the lamellipodium of mouse embryonic fibroblasts (MEFs) (6, 34). MEFs co-transfected with mEos2-fused proteins and α-actinin-GFP, as a lamellipodium reporter, were spread on glass coverslips coated with fibronectin (FN). To limit experimental variability, we performed acquisitions on cells displaying isotropic spreading which is powered by lamellipodia devoid of filopodia or mature focal adhesions (4, 5, 40). Since Rac1 is activated by growth factors (11), we performed all experiments in the absence of serum to study only integrin-dependent Rac1 activation (41, 42). Sequences of sptPALM were acquired (30 s at 50 Hz) in between α-actinin-GFP images to visualized lamellipodium displacements (Fig. 1, Video S1). We reconstructed and analyzed thousands of mEos2-fused proteins trajectories, sorted between the lamellipodium tip and the region outside of the lamellipodium (Fig. 1G, see methods). For trajectories lasting more than 260 ms (> 13 points), we computed the mean squared displacement (MSD), which describes the diffusive properties of a molecule. We sorted trajectories according to their diffusion modes (immobile, confined, free-diffusive) and extracted their diffusion coefficients (D) (Fig. 1, Table. S1, Video S1, see methods) (6, 34). Within the spatial resolution of our experiments (∼59 nm), all molecules with a D inferior to 0.011 μm^2^.s^-1^ are classified as immobile.

**Figure 1.**
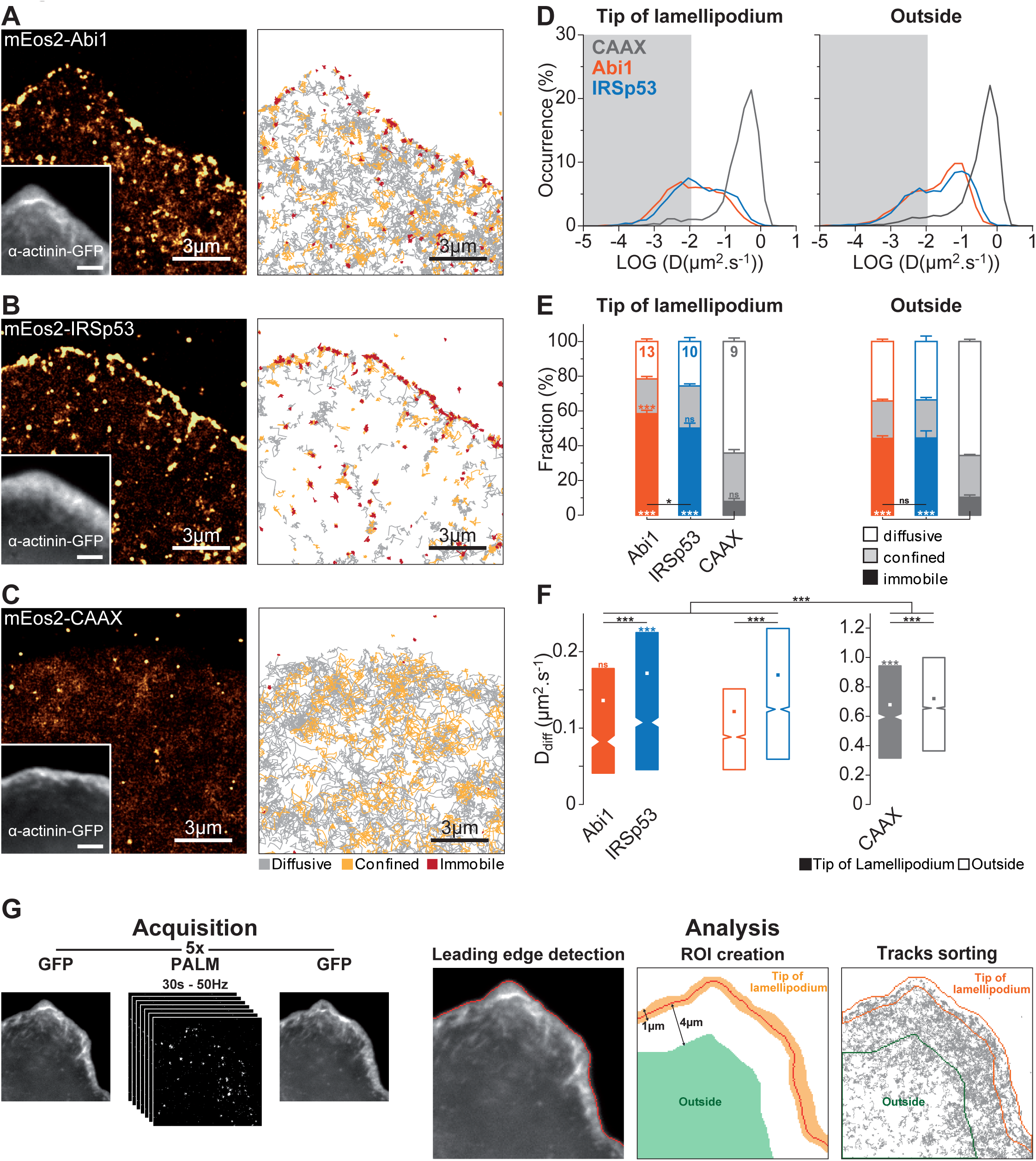
Membrane free-diffusion and trapping of the WAVE complex and IRSp53 at the lamellipodium tip. **(A)** Super–resolution intensity image of mEos2-Abi1 in the lamellipodium of a spreading MEF obtained from a sptPALM sequence (left)(50 Hz, duration: 150 s; inset: fluorescence image of α-actinin-GFP). Scale bars, 3 μm. Corresponding trajectories are color-coded to show their diffusion modes: diffusive (gray), confined (yellow) and immobile (red) (right). Scale bars, 3 μm. **(B)** Same as A for mEos2-IRSp53. **(C)** Same as A for mEos2-CAAX. **(D)** Distribution of LOG(D) for mEos2-Abi1 (orange), mEos2-IRSp53 (blue), and mEos-CAAX (gray) at the lamellipodium tip (left) and outside the lamellipodium (right; mean for cells). The grey areas including D values inferior to 0.011 μm^2^.s^-1^ correspond to immobile trajectories. **(E)** Fraction of diffusive, confined and immobile populations at the lamellipodium tip (left) and outside the lamellipodium (right), mean ± SEM for cells. **(F)** Diffusion coefficient (D) for free-diffusive trajectories at the lamellipodium tip (left) and outside the lamellipodium (right) were represented by box plots displaying the median (notch) and mean (square) ± percentile (25-75%). **(G)** Schematic representation of the experimental and analysis work flow (see methods). Sequences of sptPALM were acquired (5 x 1500 images at 50 Hz) in between α-actinin-GFP images to localize the lamellipodium (left). The cell edge was detected for each α-actinin-GFP images to define the region corresponding to the lamellipodium tip. The region outside the lamellipodium was defined as the region 4 μm inward from the lamellipodium tip. Trajectories were sorted between the lamellipodium tip and the region outside of the lamellipodium. All results for each condition correspond to pooled data from several independent experiments (cells/trajectories): Abi1 (13/8,982), IRSp53 (10/11,589) and CAAX (9/29,938), also see Table S1. Where indicated, statistical significances were obtained using two-tailed unpaired t-test for fractions of immobilization (E) or non-parametric, two-tailed Mann–Whitney rank sum test for diffusion coefficient (F). For lamellipodium tip and outside the lamellipodium, the different conditions were compared (a black line indicates which conditions were compared). For each condition fractions of immobilization (E) and diffusion coefficient (F) at lamellipodium tip and outside the lamellipodium were compared (P-values were indicated by a colored information). The resulting P-values are indicated as follows: ns, P > 0.05; *P < 0.05; ***P < 0.001.

To characterize the diffusive behavior of a control protein anchored to the inner leaflet of the plasma membrane, we used mEos2-CAAX, which displayed low fraction of immobilization and fast free-diffusion both at the lamellipodium tip (8 ± 2 %; D_diff_ = 0.678 μm^2^.s^-1^) and at the membrane outside the lamellipodium (10 ± 1 %; D_diff_ = 0.721 μm^2^.s^-1^) (Fig. 1C, D-F, Table S1). From sptPALM sequences, we generated single-molecule-based super-resolution intensity images (28, 43), showing no selective immobilizations of mEos2-CAAX at the lamellipodium tip (Fig. 1C). Actin nucleation events driven by the WAVE complex are localized at the plasma membrane of the lamellipodium tip (15, 44, 45). The WAVE complex is stable and composed of five subunits: Wave2, Abi1, Nap1, Brick and Sra (46, 47). The WAVE complex subunit, mEos2-Abi1, displayed a larger fraction of membrane free-diffusion outside the lamellipodium compared to the lamellipodium tip (tip: 22 ± 2 %; outside: 34 ± 2 %)(1A, D-F, Video S1, Table S1). Super-resolution intensity images showed the selective immobilizations of mEos2-Abi1 at the lamellipodium tip (Fig. 1A), corresponding to locations where repetitive mEos2 fluorescence signals were detected, and explained the increased fraction of immobilization at this location (tip: 58 ± 2 %; outside: 44 ± 2 %)(Fig. 1D,E, Video S1). Thus our results suggest that the WAVE complex is recruited to the lamellipodium tip by a diffusion trapping mechanism, in line with a previous study using SPT in *Xenopus* tissue culture cells (48). Those selective immobilizations are not mediated by protein crowding or membrane curvature, since the diffusive behavior of mEos2-CAAX was identical inside and outside the lamellipodium tip, but probably reflect specific interactions with proteins at the lamellipodium tip. We next examined the behavior of a I-BAR domain protein, IRSp53, which drives negative membrane curvature (49) and has been involved in Rac signaling to WAVE (15, 50–52). The diffusive behavior of mEos2-IRSp53 is similar to mEos2-Abi1, exhibiting membrane free-diffusion out of the lamellipodium and specific immobilization at the lamellipodium tip (Fig. 1B, D-F, Table S1). Thus like WAVE, IRSp53 is recruited at the lamellipodium tip by a diffusion-trapping mechanism.

### Rac1 immobilization at the lamellipodium tip is correlated with its activation state

Many studies demonstrated that the WAVE complex is a critical downstream target of Rac1 triggering cell edge protrusion (11–13). To test if selective immobilizations of the WAVE complex were triggered by interactions with Rac1 bound to the lamellipodium tip we performed sptPALM acquisition on wild type (WT) Rac1. The molecular dynamics of mEos2-Rac1-WT was largely dominated by membrane free-diffusion at the lamellipodium tip and at the membrane outside the lamellipodium (tip: 58 ± 2 %; outside: 62 ± 1 %) (Fig. 2A, D-F, Table S1). Although the fraction of immobilization was not significantly increased at the lamellipodium tip compared to CAAX-mEos2 (Rac1: 13 ± 2 %; CAAX: 8 ± 2 %) (Fig. 2E), the mean coefficient of free-diffusion (D_diff_) was significantly decreased (Rac1: D_diff_ = 0.482 μm^2^.s^-1^; CAAX: D_diff_ = 0.678 μm^2^.s^-1^) (Fig. 2F), suggesting that Rac1 transiently interacts with biomolecules at the lamellipodium tip, slowing down its movements. These results demonstrated that Rac1 is not responsible for the selective immobilization of WAVE at the lamellipodium tip.

**Figure 2.**
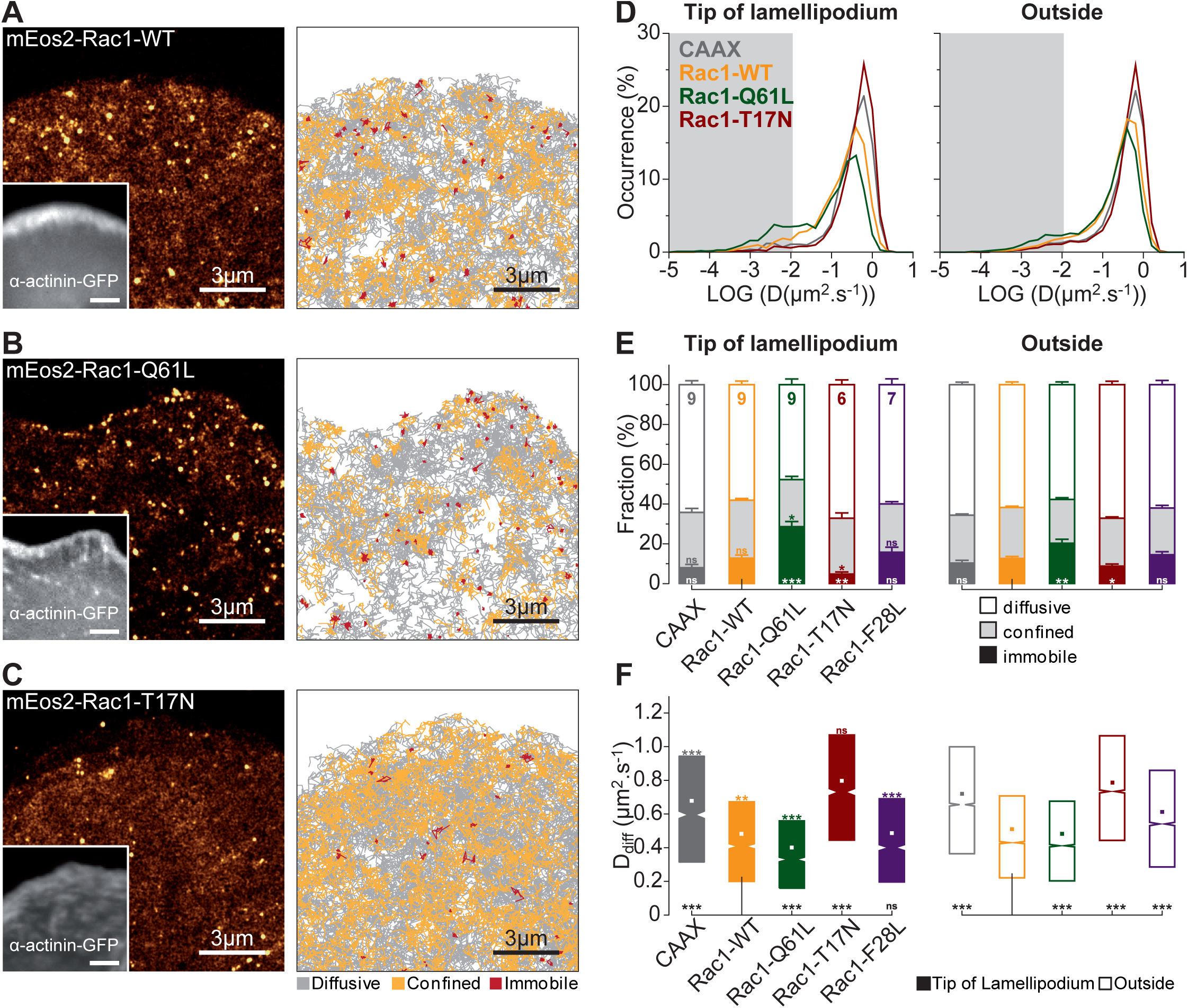
Correlation between Rac1 activation level and transient immobilizations at the LM tip. **(A)** Super–resolution intensity image of mEos2-Rac1-WT in the lamellipodium of a spreading MEF obtained from a sptPALM sequence (left)(50 Hz, duration: 150 s; inset: fluorescence image of α-actinin-GFP). Scale bars, 3 μm. Corresponding trajectories are color-coded to show their diffusion modes: diffusive (gray), confined (yellow) and immobile (red) (right). Scale bars, 3 μm. **(B)** Same as A for mEos2-Rac1-Q61L. **(C)** Same as A for mEos2-Rac1-T17N. **(D)** Distribution of LOG(D) for mEos2-Rac1-WT (yellow), mEos2-Rac1-Q61L (green), mEos2-Rac1-T17N (red), and mEos-CAAX (gray) at the lamellipodium tip (left) and outside the lamellipodium (right; mean for cells). The grey areas including D values inferior to 0.011 μm^2^.s^-1^ correspond to immobile trajectories. **(E)** Fraction of diffusive, confined and immobile populations at the lamellipodium tip (left) and outside the lamellipodium (right), mean ± SEM for cells. mEos2-Rac1-WT (yellow), mEos2-Rac1-Q61L (green), mEos2-Rac1-T17N (red), mEos2-Rac1-F28L (violet), and mEos-CAAX (gray). **(G)** Diffusion coefficient for free-diffusive trajectories at the lamellipodium tip (left) and outside the lamellipodium (right) were represented by box plots displaying the median (notch) and mean (square) ± percentile (25-75%). All results for each condition correspond to pooled data from several independent experiments (cells/trajectories): Rac1-WT (9/28,852), Rac1-Q61L (9/23,068), Rac1-T17N (6/23,950), and Rac1-F28L (7/18,406), also see Table S1. Where indicated, statistical significance was obtained using two-tailed unpaired t-test for fractions of immobilization (E) or non-parametric, two-tailed Mann–Whitney rank sum test for diffusion coefficient (F). For lamellipodium tip and outside the lamellipodium, the different conditions were compared with the respective Rac1-WT condition (a black line indicates which conditions were compared). For each condition fractions of immobilization and diffusion coefficient (F) at lamellipodium tip and outside the lamellipodium were compared (P-values were indicated by a colored information). The resulting P-values are indicated as follows: ns, P > 0.05; *P < 0.05;**P < 0.01; ***P < 0.001.

A correlation between Rac1 activation and its immobilization was demonstrated in adhesive structures including focal adhesion and dendritic spines (34, 53). To determine whether Rac1 activation state controls its localization and diffusive behavior in the lamellipodium, we performed sptPALM experiments on well characterized Rac1 mutants. The constitutively active Rac1 mutant (Rac1-Q61L) is locked in its GTP-bound state which is the conformation binding and activating Rac1 targets. Strikingly, on the contrary to Rac1-WT, super-resolution intensity images demonstrated the selective immobilization of mEos2-Rac1-Q61L at the lamellipodium tip (Fig. 2B, D-E). Distributions of D were shifted towards slower diffusion (Fig. 2D), fractions of immobilization at the lamellipodium tip were increased (Rac1-Q61L: 29 ± 3 %; Rac1-WT: 13 ± 2 %), but did not reached the level of immobilization measured for the WAVE complex (58 ± 2 %). These results suggested that Rac1-Q61L immobilization at the lamellipodium tip are transient events and could explain why classical fluorescence microscopy could not reveal enrichment of active Rac1 at the lamellipodium tip (42). In agreement with increased interactions of Rac1-Q61L with its targets, D_diff_ are also significantly decreased at the lamellipodium tip compared to Rac1-WT (Rac1-Q61L: D_diff_ = 0.401 μm^2^.s^-1^; Rac1-WT: D_diff_ = 0.482 μm^2^.s^-1^) (Fig. 2F). Interestingly, mEos2-Rac1-F28L fast cycling mutant (54), which rapidly switches between activated (GTP-bound) and inactivated (GDP-bound) states, exhibited fraction of immobilization and diffusive behavior closer to mEos2-Rac1-WT than to mEos2-Rac1-Q61L (Fig. 2E-F, Fig. S1), indicating that Rac1-WT is either rarely active or also cycles rapidly between activation and inactivation via the action of GAPs and GEFs (55). The constitutively inactive Rac1 mutant (Rac1-T17N) is locked in its GDP-bound state which is the inactive conformation unable to bind and activate targets. Like the mEos2-CAAX membrane control, mEos2-Rac1-T17N is dominated by fast free-diffusion and does not display selective immobilization at the lamellipodium tip or decreased free-diffusion such as WT and active mutants of Rac1 (Fig. 2C, D-F). Our results demonstrated a correlation between Rac1 activation and its selective immobilization at the lamellipodium tip where the WAVE complex is located.

### RhoA does not display selective immobilization at the lamellipodium tip

FRET-based sensors were used to study the spatiotemporal pattern of Rho GTPases (Rac1, RhoA, Cdc42) activation in the lamellipodium (37, 38, 42), in conjunction with periods of lamellipodium protrusions and retractions (37). The latter study showed that Rac1 activation was the strongest at the back of the lamellipodium and that maximal Rac1 activation was not synchronized with the onset of lamellipodium protrusion. Instead, RhoA strongest activation encompassed the lamellipodium tip (38), and the activation peak was correlated with the onset of lamellipodium protrusion (37). To investigate if RhoA was selectively immobilized at the lamellipodium tip we performed sptPALM experiments using mEos2-RhoA-WT. Like Rac1-WT, RhoA-WT dynamic behavior was dominated by membrane free-diffusion (tip: 58 ± 4 %; outside: 59 ± 3 %) (Fig. 3A, D-F, Table S1). The fraction of immobilization at the lamellipodium tip and outside the lamellipodium were not significantly different (tip: 16 ± 3 %; outside: 16 ± 3 %) (Fig. 3E). Next we tested, if like Rac1, the activation state of RhoA was correlated with its selective immobilization in distinct regions of the lamellipodium. We applied the same strategy, using constitutively active (RhoA-Q63L) and inactive (RhoA-G17A) RhoA (56). On the contrary to Rac1-Q61L, active mEos2-RhoA-Q63L did not display selective immobilization at the lamellipodium tip (tip: 27 ± 3 %; outside: 27 ± 2 %) (Fig. 3B, E). This lack of selective immobilization for active RhoA was further reinforced by results obtained with inactive mEos2-RhoA-G17A, which displayed the same behavior as active RhoA-Q63L (Fig. 3C, D-F). Active and inactive RhoA mutants displayed slower free-diffusion both at the lamellipodium tip and outside compare to RhoA-WT (Fig. 3F), which suggested that they may interact more with effectors, GAPs or GEFs (55). Thus, in sharp contrast with Rac1, the activation state of RhoA does not dictate its immobilization at the lamellipodium tip.

**Figure 3.**
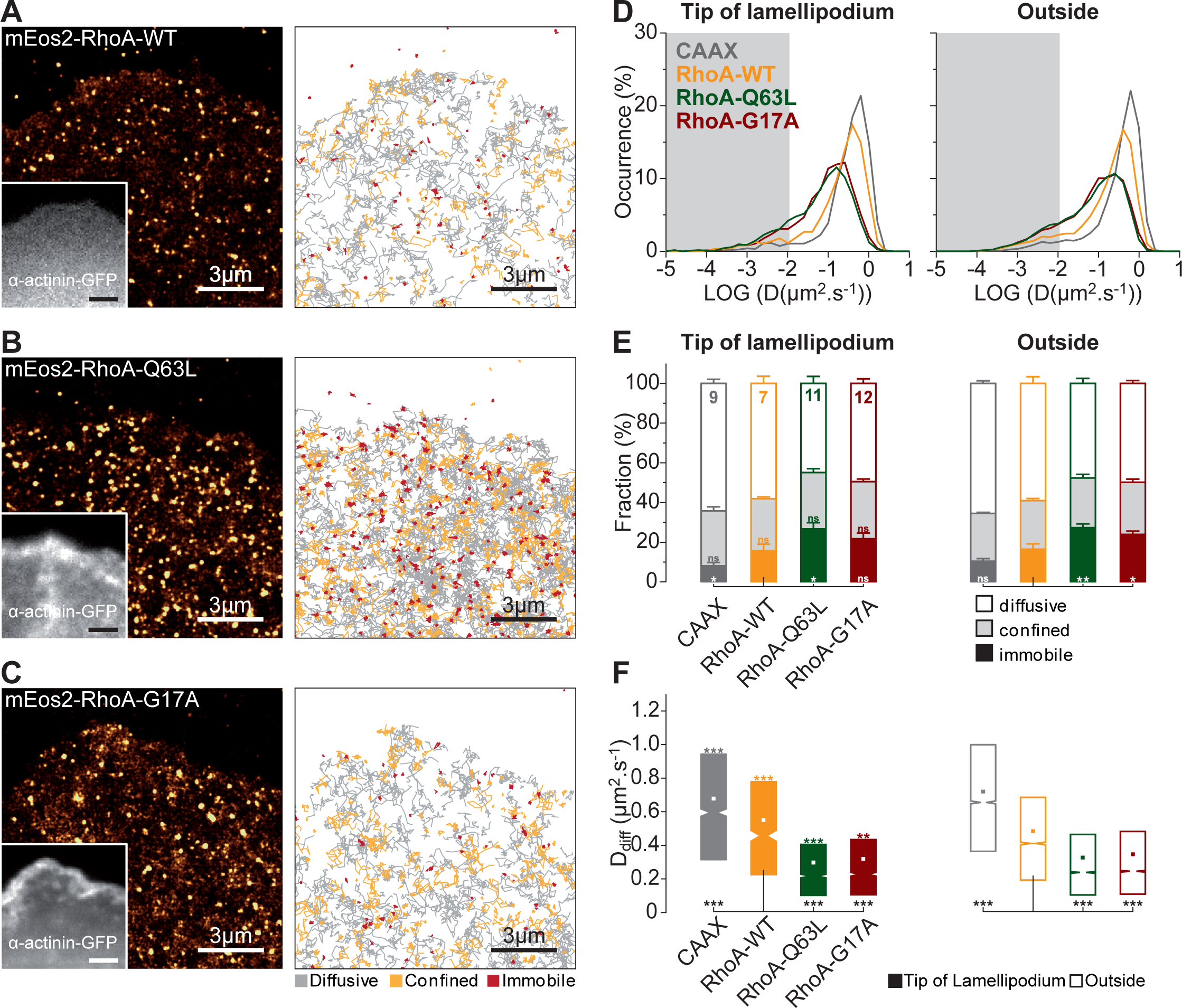
RhoA activation level is not correlated with immobilizations in specific regions of the lamellipodium. **(A)** Super–resolution intensity image of mEos2-RhoA-WT in the lamellipodium of a spreading MEF obtained from a sptPALM sequence (left)(50 Hz, duration: 150 s; inset: fluorescence image of α-actinin-GFP). Scale bars, 3 μm. Corresponding trajectories are color-coded to show their diffusion modes: diffusive (gray), confined (yellow) and immobile (red) (right). Scale bars, 3 μm. **(B)** Same as A for mEos2-RhoA-Q63L. **(C)** Same as A for mEos2-RhoA-G17A. **(D)** Distribution of LOG(D) for mEos2-RhoA-WT (yellow), mEos2-RhoA-Q63L (green), mEos2-RhoA-G17A (red), and mEos-CAAX (gray) at the lamellipodium tip (left) and outside the lamellipodium (right; mean for cells). The grey areas including D values inferior to 0.011 μm^2^.s^-1^ correspond to immobile trajectories. **(E)** Fraction of diffusive, confined and immobile populations at the lamellipodium tip (left) and outside the lamellipodium (right), mean ± SEM for cells. **(G)** Diffusion coefficient (D) for free-diffusive trajectories at the lamellipodium tip (left) and outside the lamellipodium (right) were represented by box plots displaying the median (notch) and mean (square) ± percentile (25-75%). All results for each condition correspond to pooled data from several independent experiments (cells/trajectories): RhoA-WT (7/14,817), RhoA-Q63L (11/32,839) and RhoA-G17A (12/34,069) also see Table S1. Where indicated, statistical significance was obtained using two-tailed unpaired t-test for fractions of immobilization (E) or non-parametric, two-tailed Mann–Whitney rank sum test for diffusion coefficient (F). For lamellipodium tip and outside the lamellipodium, the different conditions were compared with the respective RhoA-WT condition (a black line indicates which conditions were compared). For each condition fractions of immobilization (E) and diffusion coefficient (F) at lamellipodium tip and outside the lamellipodium were compared (P-values were indicated by a colored information). The resulting P-values are indicated as follows: ns, P > 0.05; *P < 0.05;**P < 0.01; ***P < 0.001.

### Rac1 displays fast cycles of activation and inactivation close to the lamellipodium tip

The small fraction of Rac1-WT immobilization at the lamellipodium tip compared to activated Rac1-Q61L indicated either that Rac1 activations are rare events or that the dwell time of interactions between activated Rac1 and WAVE are shorter than the acquisition frequency. To differentiate those two possibilities, we increased the acquisition frequency of sptPALM experiments to capture more efficiently transient Rac1-WT immobilizations. We switched from 50 Hz acquisition frequency using EMCCD to 333 Hz using sCMOS camera (57). The strong correlation between Rac1 activation and its selective immobilization at the lamellipodium tip was still observed at 333 Hz (Fig. 4, Fig. S2, Table S2). Moreover, increasing the acquisition frequency enhanced the differential between Rac1-WT and active Rac1 mutants (Rac1-Q61L and Rac1-F28L) (Fig. 4, Video S2-S3). Indeed, the diffusive behavior of the fast cycling Rac1-F28L mutant, that was identical to Rac1-WT at 50 Hz (Fig. 2), shifted towards the diffusive behavior of Rac1-Q61L at 333 Hz (Fig. 4). Although, we could observe seldom diffusion-trapping events of Rac1-WT at the lamellipodium tip (Video S2), this frequency was not sufficient to readily acquire the majority of Rac1-WT immobilizations that were transient, suggesting that most Rac1 interactions with the WAVE complex are transient events lasting less than 10 ms.

**Figure 4.**
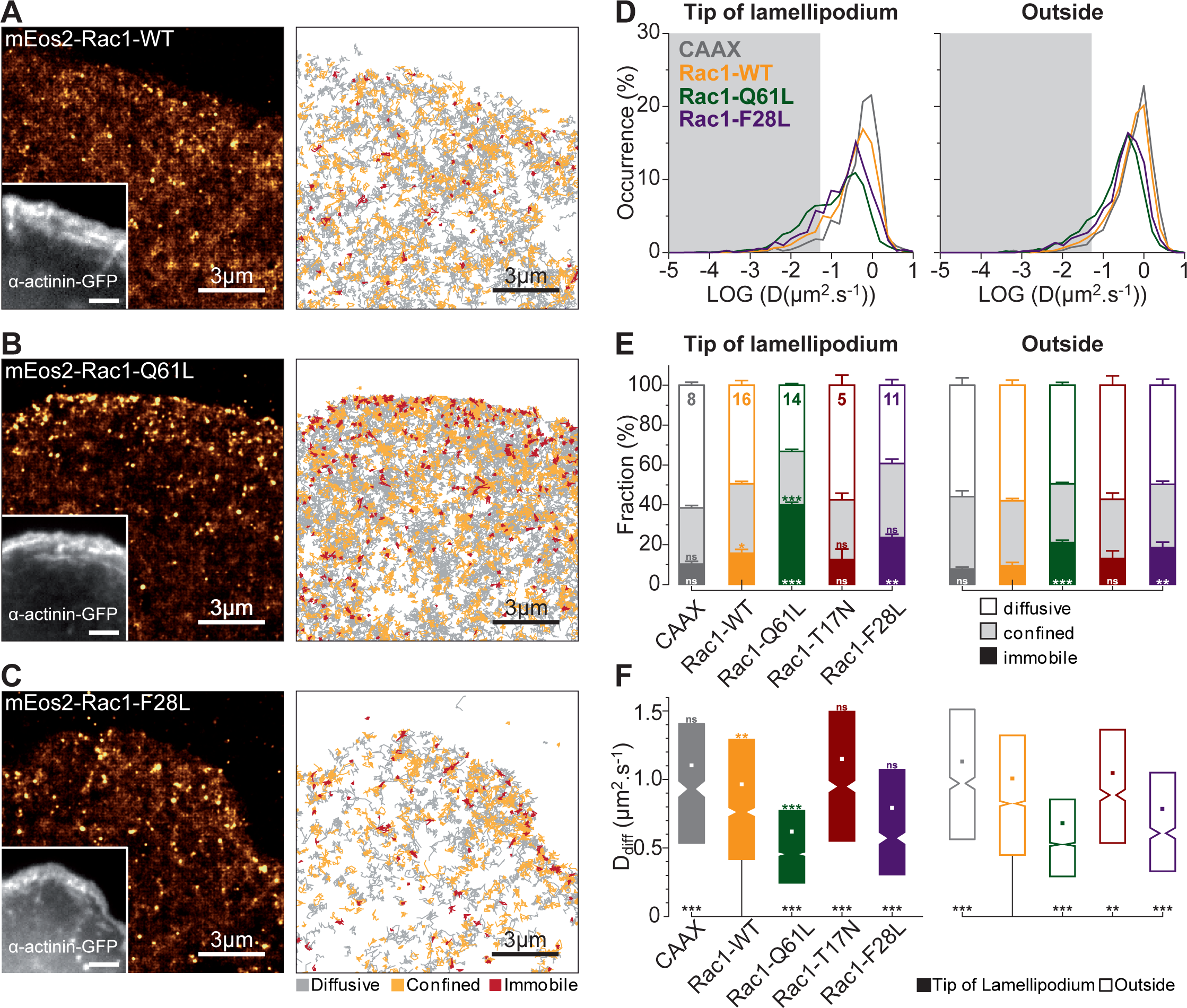
Distinct diffusive behaviors of Rac1-WT, Rac1-F28L and Rac1-Q61L revealed by fast acquisitions frequencies. **(A)** Super–resolution intensity image of mEos2-Rac1-WT in the lamellipodium of a spreading MEF obtained from a sptPALM sequence (left)(333 Hz, duration: 150 s; inset: fluorescence image of α-actinin-GFP). Scale bars, 3 μm. Corresponding trajectories are color-coded to show their diffusion modes: diffusive (gray), confined (yellow) and immobile (red) (right). Scale bars, 3 μm. **(B)** Same as A for mEos2-Rac1-Q61L. **(C)** Same as A for mEos2-Rac1-F28L. **(D)** Distribution of LOG(D) for mEos2-Rac1-WT (yellow), mEos2-Rac1-Q61L (green), mEos2-Rac1-F28L (violet), and mEos2-CAAX (gray) at the lamellipodium tip (left) and outside the lamellipodium (right; mean for cells). The grey areas including D values inferior to 0.052 μm^2^.s^-1^ correspond to immobile trajectories. **(E)** Fraction of diffusive, confined and immobile populations at the lamellipodium tip (left) and outside the lamellipodium (right), mean ± SEM for cells. mEos2-Rac1-WT (yellow), mEos2-Rac1-Q61L (green), mEos2-Rac1-T17N (red), mEos2-Rac1-F28L (violet), and mEos2-CAAX (gray). **(G)** Diffusion coefficient (D) for free-diffusive trajectories at the lamellipodium tip (left) and outside the lamellipodium (right) were represented by box plots displaying the median (notch) and mean (square) ± percentile (25-75%). All results for each condition correspond to pooled data from several independent experiments (cells/trajectories). CAAX (8/3,824), Rac1-WT (16/15,648), Rac1-Q61L (14/15,710), Rac1-T17N (5/3,767), and Rac1-F28L (11/4,527), also see Table S2. Where indicated, statistical significance was obtained using two-tailed unpaired t-test for fractions of immobilization (E) or non-parametric, two-tailed Mann–Whitney rank sum test for diffusion coefficient (F). For lamellipodium tip and outside the lamellipodium, the different conditions were compared with the respective Rac1-WT condition (a black line indicates which conditions were compared). For each condition fractions of immobilization (E) and diffusion coefficient (F) at lamellipodium tip and outside the lamellipodium were compared (P-values were indicated by a colored information). The resulting P-values are indicated as follows: ns, P > 0.05; *P < 0.05;**P < 0.01; ***P < 0.001.

To test the dependence between the location of Rac1 activation and efficient stimulation of membrane protrusion, we used local optogenetic activation of Rac1-WT. Light induced interaction of the Cryptochrome CRY2 to its membrane-anchored CIBN partner, was used to control spatially and temporally the membrane recruitment of cytosolic Rho GEFs (58). We applied the same strategy for Tiam1, a Rac1 GEF which also interacts with IRSp53 to promote WAVE activation (50).

We used the optogenetic system composed of CIBN-GFP-CAAX localized at the cell membrane and CRY2PHR-Tiam1-iRFP (Tiam1-CRY2), which is initially cytoplasmic (Fig. 5A) (Remorino et al., companion manuscript). Blue light illumination triggered the fast and local membrane recruitment of Tiam1, visualized by enrichment of the Tiam1-CRY2 fluorescent signal (Fig. 5B,C, Video S4). We performed the experiments on already formed lamellipodium, illuminating regions at the lamellipodium back or encompassing the lamellipodium tip (Fig. 5D). When the region enclosed the lamellipodium tip, membrane recruitment of Tiam1-CRY2 rapidly increased the speed of protrusion (Fig. 5E, Video S4). Importantly, stimulation of membrane protrusion was inefficient when the illuminated region was located ∼3 μm away from the tip (Fig. 5E), despite efficient Tiam1-CRY2 membrane translocation. These results suggest that Rac1 activation must occur close to its targets at the lamellipodium tip, and that inactivation of Rac1 arises rapidly out of the illuminated region.

**Figure 5.**
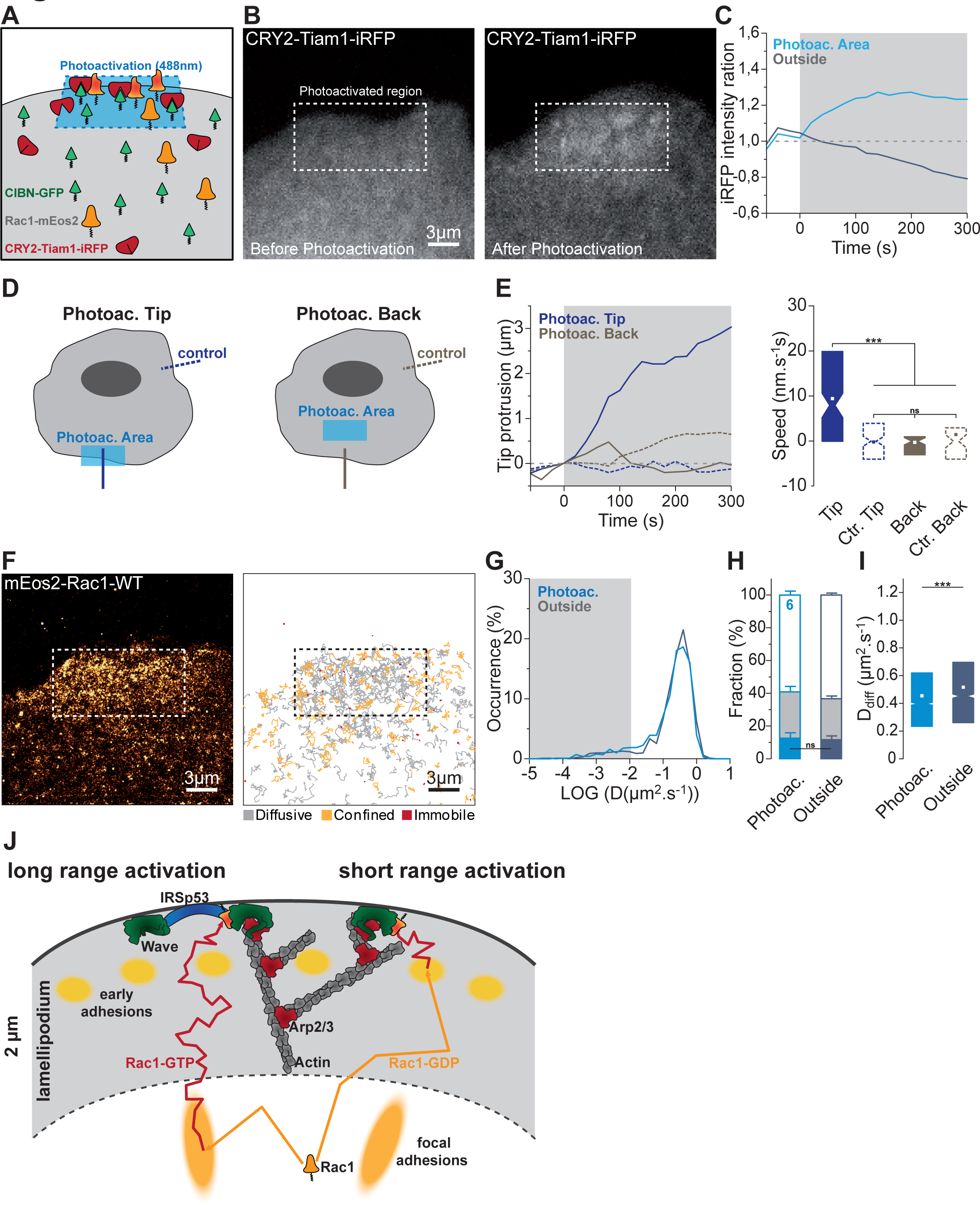
Rac1 must be activated at the lamellipodium tip to stimulate membrane protrusion. **(A)** Schematic description of Rac1 optogenetic activation using CIBN-GFP-CAAX localized at the cell membrane, cytosolic CRY2PHR-Tiam-iRFP (Tiam-CRY2). Local photoactivation using a 488 nm laser induces CRY2PHR-Tiam-iRFP membrane recruitment triggering Rac1 enhanced activation (Video S4). **(B)** TIRF images of CRY2PHR-Tiam-iRFP before photocativation (left), after photoactivation (right). **(C)** Normalized fold increase of CRY2PHR-Tiam-iRFP fluorescence, acquired in the TIRF mode, in the photoactivated region (blue) or outside (gray) before and during photoactivation (gray area). **(D)** Schematic representation of photoactivation areas. Photoactivation encompassing the lamellipodium tip (Photoac. Tip) and 3 μm away from the lamellipodium tip (Photoac. Back). **(E)** lamellipodium protrusion as function of time (left) before and during photoactivation (gray area) and protrusion speed during photoactivation (right) for the area encompassing the lamellipodium tip (dark blue) and the area away from the lamellipodium tip (gray). Protrusions were measured in front of photoactivation areas (plain lines) and in control parts of the cells (dotted lines), as schematized in (D). **(F)** Super–resolution intensity image obtained from a sptPALM sequence of mEos2-Rac1-WT in the lamellipodium of a spreading MEF after optogenetic membrane recruitment of CRY2PHR-Tiam-iRFP (left)(same cell as B, 50 Hz, duration: 10 s). Scale bars, 3 μm. Corresponding trajectories are color-coded to show their diffusion modes: diffusive (gray), confined (yellow) and immobile (red) (right). Scale bars, 3 μm. **(G)** Distribution of LOG(D) for mEos2-Rac1-WT inside (blue) and outside (gray) the photoactivated region of interest. The grey areas including D values inferior to 0.011 μm^2^.s^-1^ correspond to immobile trajectories. **(H)** Fraction of diffusive, confined and immobile populations inside (blue) and outside (gray) the photoactivated region of interest. **(I)** Diffusion coefficient (D) for free-diffusive trajectories inside (blue) and outside (gray) the photoactivated region of interest were represented by box plots displaying the median (notch) and mean (square) ± percentile (25-75%). All results for each condition correspond to pooled data from several independent experiments (cells/trajectories); Rac1-WT (6/5,689) also see Table S1. **(J)** Schematic representation of the working model. Where indicated, statistical significance was obtained using two-tailed unpaired t-test for fractions of immobilization (H) or non-parametric, two-tailed Mann– Whitney rank sum test for diffusion coefficient (I) and protrusion speed (E), the resulting P-values are indicated as follows: ns, P > 0.05; *P < 0.05;**P < 0.01; ***P < 0.001.

To prove further the passing nature of Rac1 and WAVE interactions, independently from the use of constitutively active mutants, we combined optogenetic activation of Rac1-WT with simultaneous tracking of mEos2-Rac1-WT (Fig. 5F-I). Sequences of mEos2-Rac1-WT sptPALM (30 s, 50 Hz) were acquired in between blue light illumination of a region enclosing the lamellipodium tip (Fig. 5F). Membrane re-localization of Tiam-CRY2, immediately increased the speed of membrane protrusion, accounting for the local enhanced activation of Rac1-WT. Nevertheless, the diffusive properties of Rac1-WT inside and outside the region of optogenetic activation were similar (Fig. 5G-I, Table S1). Altogether, these results suggested that Rac1 activation events, which are associated with Rac1 immobilizations, are transient and local events.

## Discussion

Combining SPT with loss- or gain-of-function of Rho GTPases mutants, we linked the nanoscale dynamic localization of Rac1 to its activation and site of action within the lamellipodium. We showed that the Rac1 effector WAVE and Rac1 regulator IRSp53 accumulate at the lamellipodium tip by membrane free-diffusion and trapping. We demonstrated a strong correlation between Rac1 activation state and its immobilization at the lamellipodium tip. In contrast, RhoA activation state is not correlated with increased immobilizations within specific regions of the lamellipodium. Those results suggest that Rac1 action during lamellipodium formation occurs primarily at the lamellipodium tip. Despite direct interactions with WAVE and IRSp53 (52), Rac1-WT only displays a decreased diffusion at the lamellipodium tip, suggesting fast and local activation/inactivation cycles. Indeed, coupling sptPALM with optogenetic activation of Rac1, triggered by Tiam1 membrane recruitment, demonstrates that Rac1-WT diffusive properties are unchanged despite enhanced lamellipodium protrusion. Furthermore, local optogenetic Rac1 activation demonstrates that Rac1 must be activated at the lamellipodium tip and not at the lamellipodium back to stimulate protrusion. Thus, our data show that Rac1 is rapidly cycling between activation and inhibition at the proximity of its targets, ensuring a local and fast control over Rac1 actions.

### The site of Rac1 activation can differ from its site of action

The tight spatiotemporal regulation of actin regulatory proteins gives rise to complex periodic patterns of lamellipodium protrusions and retractions (4, 24). In the signaling model, based on protein activity measurements, the formation of periodic patterns results from a complex interplay between signaling events (37, 59). In the mechanical model, based on forces generated by the lamellipodium, it is the periodic physical connection of the lamellipodium to myosin and early adhesions that drives periodic contractions (4, 5, 60, 61). The use of Rho GTPases FRET sensors timed on periods of cell edge protrusions and retractions, enabled to correlate their signaling activities with the start or end of cell edge movements. The zone of maximal Rac1 activation was located at the lamellipodium back and found to climax asynchronously with the onset of edge protrusion (37). On the contrary, the zone of RhoA activation encompassed the cell edge and was synchronized with the onset of protrusion (38, 62). These results suggest that RhoA, potentially via formins (63, 64) could trigger lamellipodium formation, while Rac1 is not the primary signal triggering edge protrusion but could be involved in the stabilization of newly expanded protrusions. Nevertheless, our results strongly support a model where Rac1 is acting at the lamellipodium tip to trigger membrane protrusion. One way to reunite those seemingly contradictory findings is to consider that sites of protein activation could be distinct from sites of action. Signaling events propagate within a cell, enabling a local signal to have remote consequences. In addition, these conclusions were based on conventional fluorescence microscopy heat maps of Rho GTPases activation. In fact, the area of Rac1 activation, though not maximal at the lamellipodium tip, is still elevated at this location and could trigger WAVE activation (37, 42). The broad area of RhoA activation encompasses entirely the lamellipodium (37, 38), thus it is still difficult to conclude whether RhoA is acting at the lamellipodium tip or back. In agreement with a crucial role of Rac1 during cell edge protrusion, PDGF-evoked fast membrane protrusions are associated with shut down of RhoA activity at the cell periphery, while Rac1 activity concentrates at the cell periphery (65).

### Correlation between Rac1 activation and immobilization at the lamellipodium tip

All proteins are part of interaction hubs, implying that their functions depend on binding events with members of this hub. Thus, interactions should translate into changes in molecular dynamics, both for structural and signaling proteins. Using sptPALM, we demonstrated that integrin activation events are correlated with their immobilization triggered by interactions with intracellular activators or/and extracellular ligands (6, 66). However, in the case of signaling proteins, the corollary could be more difficult to prove experimentally, especially if interactions occur at time-scales well below the temporal resolution or if the binding pair moves with the same behavior as individual proteins (67). Rho GTPases are enrolled in complex signaling networks composed of myriads of GEFs, GAPs, regulators and effectors (55). The use of loss- or gain-of-function mutants could prolong interaction times with effectors or regulators above the temporal resolution, enabling to capture binding events associated with protein function. Using this strategy, previous studies demonstrated that Rac1 activation is correlated with immobilization in integrin-dependent focal adhesions (53) or neuronal dendritic spines (34). Using conventional fluorescence microscopy, constitutively active Rac1-G12V GFP fusion proteins displayed a homogeneous distribution without clear specific localization within the lamellipodium (42). SPT and super-resolution microscopy enabled us to demonstrate that active Rac1-Q61L is specifically immobilized at the lamellipodium tip, like the WAVE complex and IRSp53. Likewise, in the companion manuscript, the level of Rac1 activation was correlated with Rac1 immobilization in signaling nanoclusters in protrusive region of polarized cells (Remorino et al.). Nevertheless, the constitutively inactive Rac1-T17N mutant and the CAAX-polybasic region were also specifically immobilized in signaling nanocluster. On the contrary, neither Rac1-T17N nor CAAX exhibited significant immobilizations at the lamellipodium tip, suggesting that the signaling nanoclusters described in the companion manuscript are not localized at the lamellipodium tip in migrating cells. The fast cycling mutation Rac1-F28L, which is comparable to the Rac1-P29S mutation found in melanoma (54, 68), display increased fraction of immobilization and decreased rate of free-diffusion, compared to Rac1-WT at 333 Hz, indicating an increased binding activity towards its effectors. On the contrary to Rac1-Q61L, this increased activity was not sufficient to induce a significant Rac1-F28L enrichment at the lamellipodium tip. Despite the potential role of RhoA in triggering the onset of membrane protrusions (37) through formin-dependent F-actin nucleation and elongation (64, 69), no specific immobilization zones were detected in the lamellipodium. The binding partner responsible for active Rac1 immobilization at the lamellipodium tip is unknown. Integrins could induce local Rac activation and binding with its effectors by directing Rac membrane recruitment through dissociation from Rho-GDI (42, 70). Furthermore, Rac1-WT and constitutively active Rac1 are immobilized in mature adhesive structures (53). This is consistent with the localization of Rac1 GEFs in mature but also in early integrin-dependent adhesive structures (53, 71, 72). Thus, immobilization of active Rac1 at the lamellipodium tip could be mediated by recruitment to early adhesive structures. Nevertheless, early adhesive structures are not initiated exactly at the lamellipodium tip, but ∼400 nm back from it (5). The constitutively inactive Rac1-T17N mutant, which displays increased binding activity towards GEFs but cannot bind effectors or GAPs, behaves like the CAAX control exhibiting no immobilizations at the lamellipodium tip, which suggests that Rac1 GEFs are not concentrated at this location. Thus, active Rac1 probably interacts with protein of the lamellipodium tip such as WAVE or IRSp53 (14, 15, 52).

### Long or short range of Rac1 action

The efficiency of Rac1 action on its targets after activation, i.e. GTP loading, is timed by the rate of GTP hydrolysis, but also by the relative distance between sites of Rac1 activation and effectors. This implies that the rate of Rac1 diffusion in the cytosol or at the plasma membrane is a crucial parameter in defining the radius of Rac1 action. A recent study using SPT showed that Rac1 membrane recruitment precedes GTP loading (73). This first step is regulated by Rho GDI which sequesters Rho GTPases in the cytosol by preventing prenylated and palmitoylated CAAX tails from binding to the membrane (74). This suggests that Rac1 reaches its targets by membrane diffusion and not cytosolic diffusion. Our study demonstrates that the activation state of Rac1 will determine its diffusive properties at the lamellipodium tip, but also at the plasma membrane outside the lamellipodium. Thus, the radius of Rac1 action will be set by its dwell time of activation, consequently activated Rac1 will explore a smaller area than inactivated Rac1.

WAVE activation at the lamellipodium tip could be triggered by Rac1 activated at the lamellipodium back, or locale Rac1 activation close to the lamellipodium tip. In the long range model (Fig. 5J), Rac1 activation could occur remotely at mature adhesion sites located behind the lamellipodium, thus activated GTP-bound Rac1 must travel at least 2 μm to reach its target (37). Since the median free-diffusion of Rac1-Q61L is 0.52 μm^2^.s^-1^, Rac1-WT must remain active for few seconds to enable half of the activated Rac1 to reach the lamellipodium tip, well above our acquisition frequency (50-333 Hz). If this was the case we should have observed, for Rac1-WT, increased fraction of immobilization and shift towards lower values of Ddiff, as observed for Rac1-Q61L. Instead, all our results support the short range model, where fast Rac1 activation/inactivation cycles occur close to its location of action, the lamellipodium tip. First, forcing Rac1-WT activation using optogenetic activation at the lamellipodium tip did not increase the fraction of immobilization or decrease the Ddiff towards the levels reached by constitutively active Rac1-Q61L. Second, local optogenetic activation of Rac1 efficiently enhances protrusion if the region of activation encloses the lamellipodium tip but not at the lamellipodium back (∼3 μm). In the short range model (Fig. 5J), local Rac1 activation could be triggered by the specific localization of Rac1 GEFs either at the lamellipodium tip, like suggested by Tiam1 binding to IRSp53 (50), or in the earliest adhesive structures located less than half a micron away from the moving lamellipodium tip (5), as proposed for Tiam1 and β-Pix (71, 72).

The spatiotemporal regulation of proteins at the nanoscale is emerging as an essential feature in cell biology (8, 75, 76). Rac1 appears to display two different regimes of nanoscale dynamic organization in cells: long lasting nanoclusters, which presumably act as Rac1 signaling platforms in polarized cells (Remorino et al.); but also transient immobilizations, which are hallmarks of brief interactions with their targets during Rac1 cycling, that trigger membrane protrusions in migrating cells.

## Materials and Methods

### Cell culture and spreading assays

Mouse Embryonic Fibroblasts (MEFs) were cultured in DMEM (Gibco and Eurobio) with 10% FBS. Transient transfections of plasmids were performed 2 days before experiments using Amaxa Nucleofector (Lonza). The cells were detached with trypsin/EDTA (0.05% for 2 min), the trypsin was inactivated using soybean trypsin inhibitor (1 mg/ml in DMEM), and the cells were washed and suspended in serum-free Ringer, and incubated for 30 min before spreading on human FN (Roche)-coated glass surface (FN 10μg/ml)(4, 5).

### DNA constructs

mEos2-Abi1, mEos2-IRSp53, mEos2-CAAX, mEos2-Rac1-WT, mEos2-Rac1-Q61L, mEos2-Rac1-T17N, mEos2-Rac1-F28L were generated by PCR of the coding DNA sequence of the corresponding protein and inserted in the pcDNAm-FRT-PC-mESO2 blue at the Fse1/Asc1 sites. mEOS2-RhoA-WT, mEOS2-RhoA-Q63L and mEOS2-RhoA-G17A were generated by PCR from, respectively, pGEX-RhoA, RhoA-Q63L and RhoA-G17A (56) and cloned into the pmEOS2-C1 vector at the BglII/XhoI sites. CIBN-GFP-CAAX was described previously (58) and CRY2PHR-Tiam1-iRFP was obtained using the strategy described in (58). α-actinin-GFP construct was described previously (4). The fidelity of all constructs was verified by sequencing.

### SptPALM acquisitions

Cells were imaged at 37°C in a Ludin chamber (Life Imaging Services) with an inverted motorized microscope (Nikon Ti) equipped with a CFI Apo TIRF 100x oil, NA 1.49 objective and a perfect focus system, allowing long acquisition in TIRF illumination mode. For photoactivation localization microscopy, cells expressing mEos2 tagged constructs were photoconverted using a 405 nm laser (Omicron) and the resulting photoconverted single molecule fluorescence was excited with a 561 nm laser (Cobolt Jive^™^). Both lasers illuminated the sample simultaneously. Their respective power was adjusted to keep the number of the stochastically activated molecules constant and well separated during the acquisition. Fluorescence was collected by the combination of a dichroic and emission filters (dichroic: Di01-R561,emission: FF01-617/73, Semrock) and a sensitive EMCCD (Electron-Multiplying Charge-Coupled Device, Evolve, Photometric) for 50 Hz acquisitions or an sCMOS (scientific Complementary Metal-Oxide Semiconductor, Orca Flash4, Hamamatsu) for 333 Hz acquisitions. The acquisition was steered by Metamorph software (Molecular Devices) in streaming mode. α-actinin-GFP was imaged using a conventional GFP filter cube (excitation: FF01-472/30, dichroic: FF-495Di02, emission: FF01-520/35). Using this filter cube does not allow to separate spectrally the unconverted pool of mEos2 from the GFP fluorescent signal. However, with all the constructs used, whether the mEos2 signal was highly or poorly enriched in lamellipodium, we were still able to detect lamellipodia with α-actinin-GFP.

### Single molecule segmentation and tracking

A typical sptPALM experiment generates between 4000-7500 images per cell analysed in order to extract molecule localization and dynamics. Single molecule fluorescent spots were localized and tracked over time using a combination of wavelet segmentation and simulated annealing algorithms (77–79). Under the experimental conditions described above, the resolution of the system was quantified to 59 nm (50 Hz, EMCCD), 49 nm (333 Hz, sCMOS) (Full Width at Half Maximum, FWHM). This spatial resolution depends on the image signal to noise ratio and the segmentation algorithm (80) and was determined using fixed mEos2 samples. We analysed 2D distributions of single molecule positions belonging to long trajectories (>50 frames) by bi-dimensional Gaussian fitting, the resolution being determined as 2.3*s*_*xy*_, where *s_xy_* is the pointing accuracy.

For the trajectory analysis, lamellipodium tip were identified by thresholding the fluorescence signal of α-actinin-GFP images acquired in between sptPALM sequences (1500 images at 50 Hz and 4000 at 333 Hz). The cell edge for each α-actinin-GFP images was detected. For consecutive images, the localized edges were expended by 3 pixels (480 nm) outside of the cell for the distal edge and inside the cell for the proximal edge, delimiting the area explored by the lamellipodium tip during the intervening sptPALM acquisition. This allowed to keep the protruding lamellipodium tip in the region of interests during sptPAM sequences. The region outside the lamellipodium was defined as the region 25 pixels (4 μm) inward from the distal edge. The corresponding binary masks were used to sort single particle data analyses to specific regions, lamellipodium tip, outside the lamellipodium. We analyzed trajectories lasting at least 13 points (≥ 260 ms, 50 Hz; ≥ 39 ms, 333 Hz) with a custom routine written for Matlab using the mean squared displacement *MSD* computed as (Eq. 1):

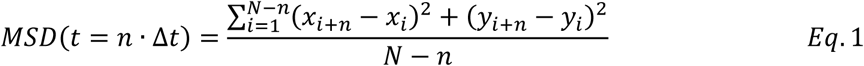

where *x*_*i*_ and *y*_*i*_ are the coordinates of the label position at time *i* · Δ*t*. We defined the measured diffusion coefficient *D* as the slope of the affine regression line fitted to the *n* = 1 to 4 values of the *MSD*(*n* · Δ*t*). The *MSD* was computed then fitted on a duration equal to 80% (minimum of 10 points, 200 ms (50 Hz), 30 ms (333 Hz)) of the whole stretch by (Eq 2):

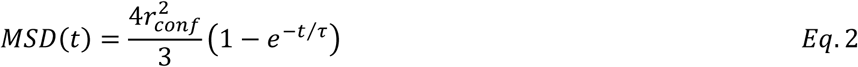

where *r*_*conf*_ is the measured confinement radius and τ the time constant 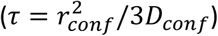. To reduce the inaccuracy of the MSD fit due to down sampling for larger time intervals, we used a weighted fit. Trajectories were sorted in 3 groups: immobile, confined diffusion and free-diffusion. Immobile trajectories were defined as trajectories with D<0.011 μm^2^.s^-1^ (50 Hz), D<0.052 μm^2^.s^-1^ (333 Hz), corresponding to molecules which explored an area inferior to the one defined by the image spatial resolution ∼(0.059 μm)^2^ at 50 Hz ∼(0.049 μm)^2^ at 333 Hz during the time used to fit the initial slope of the MSD (6). For the EMCCD experiments performed at 50 Hz, 4 points, 80 ms: D_threshold_=(0.059 μm)^2^/(4x4x0.02 s)∼0.011 μm^2^.s^-1^. For the sCMOS experiments performed at 333 Hz, 4 points, 12 ms: D_threshold_=(0.059 μm)^2^/(4x4x0.003 s)∼0.052 μm^2^.s^-1^. To separate trajectories displaying free-diffusion from confined diffusion, we used the time constant τ calculated for each trajectory. Confined and free-diffusing trajectories were defined as trajectories with a time constant τ respectively inferior and superior to half the time interval used to compute the MSD (100 ms (50 Hz); 15 ms (333 Hz)).

### Optogenetic activation of Rac1 using Tiam1 membrane recruitment

MEFs were co-transfected with CIBN-GFP-CAAX, CRY2PHR-Tiam-iRFP and mEos2-Rac1-WT. TIRF images of spreading MEFs were acquired using an azimuthal TIRF module (ilas2; Roper Scientific, Tucson, AZ) and laser power and exposure time were chosen to limit phototoxicity which could stop cell edge protrusion. To photoactivate CRY2, we used a fluorescence recovery after photobleaching (FRAP) head (Roper Scientific) to illuminate with a low 488 nm laser power (5–10%) defined regions of interest. CRY2PHR-Tiam-iRFP images were acquired every 20 s. After a base line of CRY2PHR-Tiam-iRFP images (4 images), CRY2PHR-Tiam-iRFP acquisitions were preceded by CRY2 photoactivation (∼ 500 ms). sptPALM sequences of mEos2-Rac1-WT (10 s, 500 images, 50 Hz) were acquired in between CRY2 photoactivation. This algorithm was repeated for at least 15 times.

Measurements of the lamellipodium protrusion speeds (Fig. 5E) were performed using kymographs (ImageJ plugin, Kymo ToolBox, F. Cordelières). We first generated CRY2PHR-Tiam-iRFP time-lapse sequences (Video S4). Then, Kymo ToolBox was used to generate kymographs from lines perpendicular to the lamellipodium tip in front of the photoactivated area or perpendicular to a control part of the cell (Fig. 5D). Then, lines were manually generated on kymographs, and Kymo ToolBox extracted the displacements and the speed of the cell edges.

## Acknowledgments

We thank B. Tessier, R. Sterling for technical assistance; M. Garcia, A. Gautreau, M. Lagardère, T. Orré for helpful discussions; C. Poujol, S. Marais (Bordeaux Imaging Center, BIC) for technical help; A. Gautreau and J.J. Gautier for mEos2 constructs of actin regulators. We acknowledge financial support from the French Ministry of Research and CNRS, ANR grant Integractome (GG), ANR grant FastNano (GG), Ligue Contre le Cancer (AM), Conseil Régional Aquitaine (AM), Fondation pour la Recherche Médicale (GG, AM), Fondation ARC (VM).

## Supporting Information

**Figure S1.**
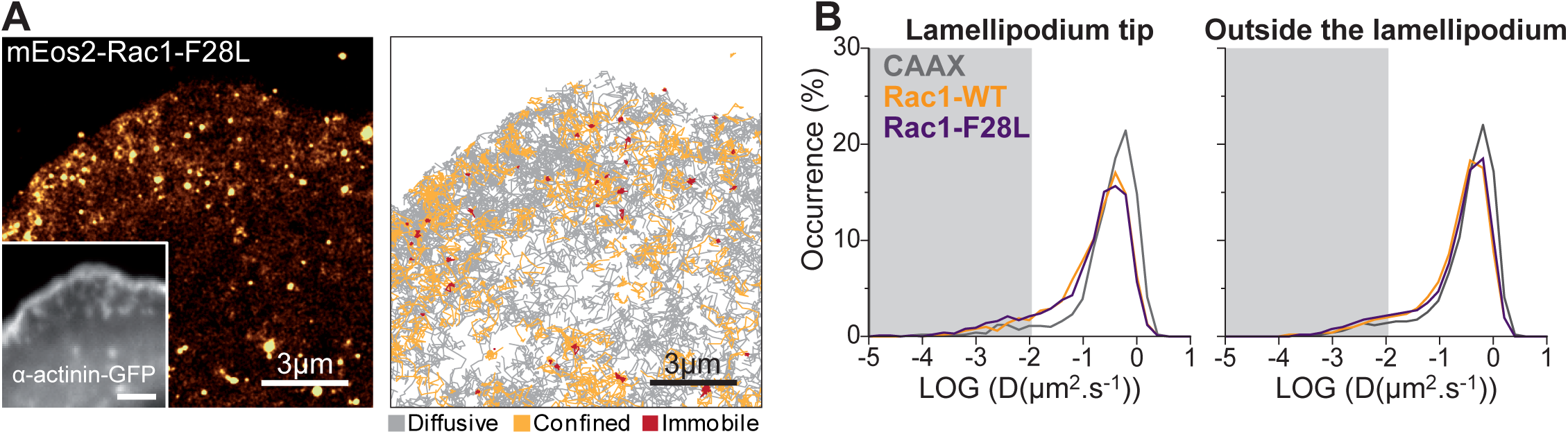
Correlation between Rac1 activation level and transient immobilizations at the LM tip. **(A)** Super–resolution intensity image of mEos2-Rac1-F28L in a spreading MEF obtained from a sptPALM sequence (left)(50 Hz, duration: 150 s; inset: fluorescence image of α-actinin-GFP). Scale bars, 3 μm. Corresponding trajectories are color-coded to show their diffusion modes: diffusive (gray), confined (yellow) and immobile (red) (right). Scale bars, 3 μm. **(B)** Distribution of LOG(D) for mEos2-Rac1-F28L (violet), mEos2-Rac1-WT (yellow), and mEos-CAAX (gray) at the lamellipodium tip (left) and outside the lamellipodium (right; mean for cells). The grey areas including D values inferior to 0.011 μm^2^.s^-1^ correspond to immobile trajectories.

**Figure S2.**
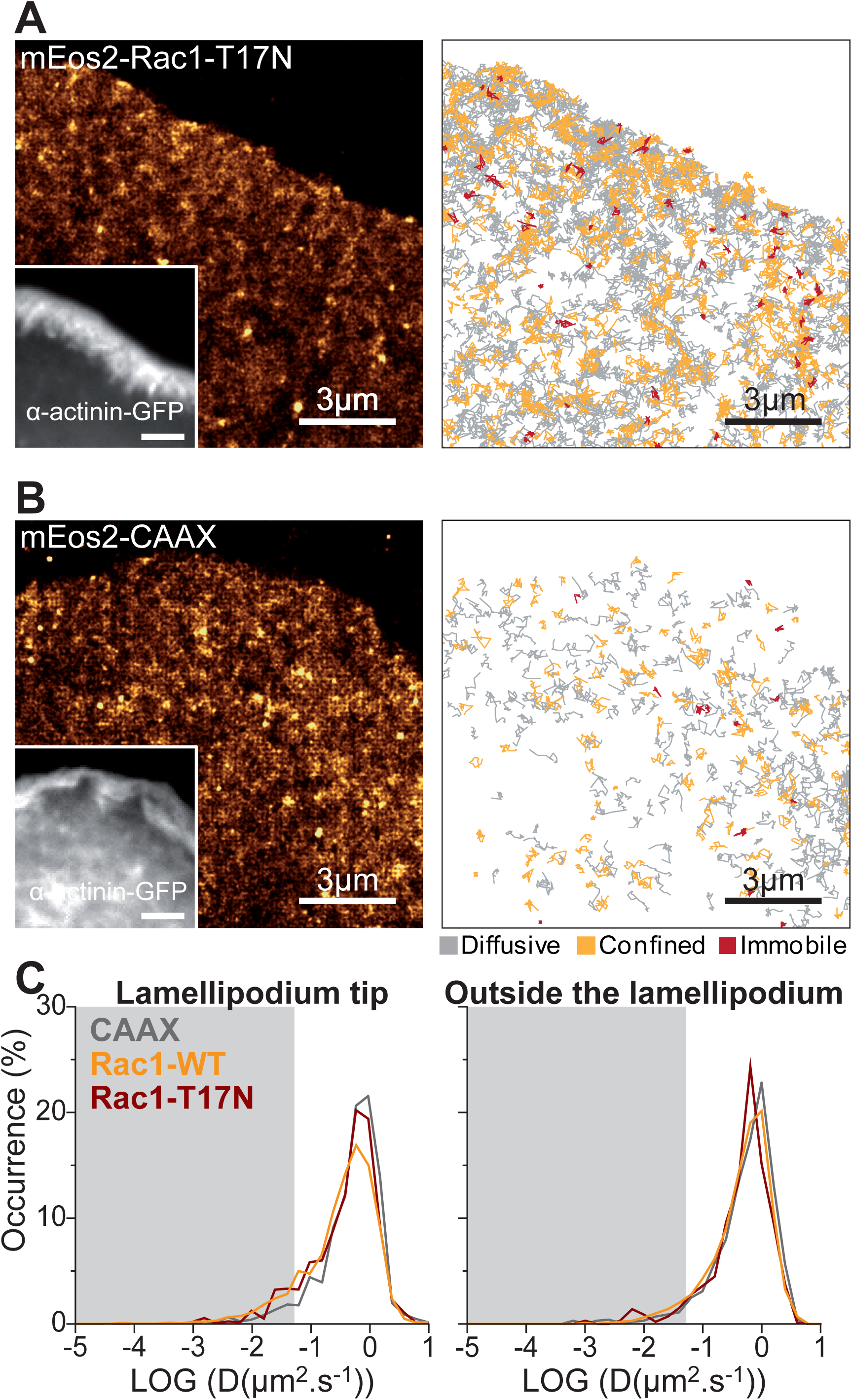
Distinct diffusive behaviors of Rac1-WT, Rac1-F28L and Rac1-Q61L revealed by fast acquisitions frequencies. **(A)** Super–resolution intensity image of mEos2-Rac1-T17N in a spreading MEF obtained from a sptPALM sequence (left)(333 Hz, duration: 150 s; inset: fluorescence image of α-actinin-GFP). Scale bars, 3 μm. Corresponding trajectories are color-coded to show their diffusion modes: diffusive (gray), confined (yellow) and immobile (red) (right). Scale bars, 3 μm. **(B)** Same as A for mEos2-CAAX. **(C)** Distribution of LOG(D) for mEos2-Rac1-T17N (red), mEos2-Rac1-WT (yellow), and mEos-CAAX (gray) at the lamellipodium tip (left) and outside the lamellipodium (right; mean for cells). The grey areas including D values inferior to 0.052 μm^2^.s^-1^ correspond to immobile trajectories.

**Video S1. From raw sptPALM acquisitions to super-resolution intensity images and trajectories.**

(**A)** Raw acquisition of mEos2-Abi1 from one sptPALM sequence (50 Hz). **(B)** Localization of single molecule fluorescent spots. **(C)** Corresponding super–resolution intensity image. **(D)** Corresponding trajectories are color-coded to show their diffusion modes: diffusive (gray), confined (yellow) and immobile (red). Scale bars, 2 μm.

**Video S2. Rare diffusion-trapping events of Rac1-WT at the LM tip.**

(**A)** Fluorescence image of α-actinin-GFP. Scale bars, 2 μm. **(B)** Raw acquisition of mEos2-Rac1-WT from one sptPALM sequence (333 Hz, duration: 1.2s). Lines were added to distinguish the lamellipodium tip. Arrow heads highlight immobilization events. Scale bars, 2 μm.

**Video S3. Diffusion-trapping events of Rac1-Q61L at the LM tip**

**(A)** Fluorescence image of α-actinin-GFP. Scale bars, 2 μm. **(B)** Raw acquisition of mEos2-Rac1-Q61L from one sptPALM sequence (333 Hz, duration: 1.2s). Lines were added to distinguish the lamellipodium tip. Arrow heads highlight immobilization events. Scale bars, 2 μm.

**Video S4. Stimulation of lamellipodium protrusion by CRY2PHR-Tiam-iRFP membrane recruitment.**

Time-lapse of CRY2PHR-Tiam-iRFP photoactivation sequences (20 s). After a base line of 4 images, a region encompassing the lamellipodium tip is photoactivated every 20 s by low 488 nm laser illumination triggering CRY2PHR-Tiam-iRFP membrane recruitment and lamellipodium protrusion. Scale bars, 15 μm

**Table S1.**
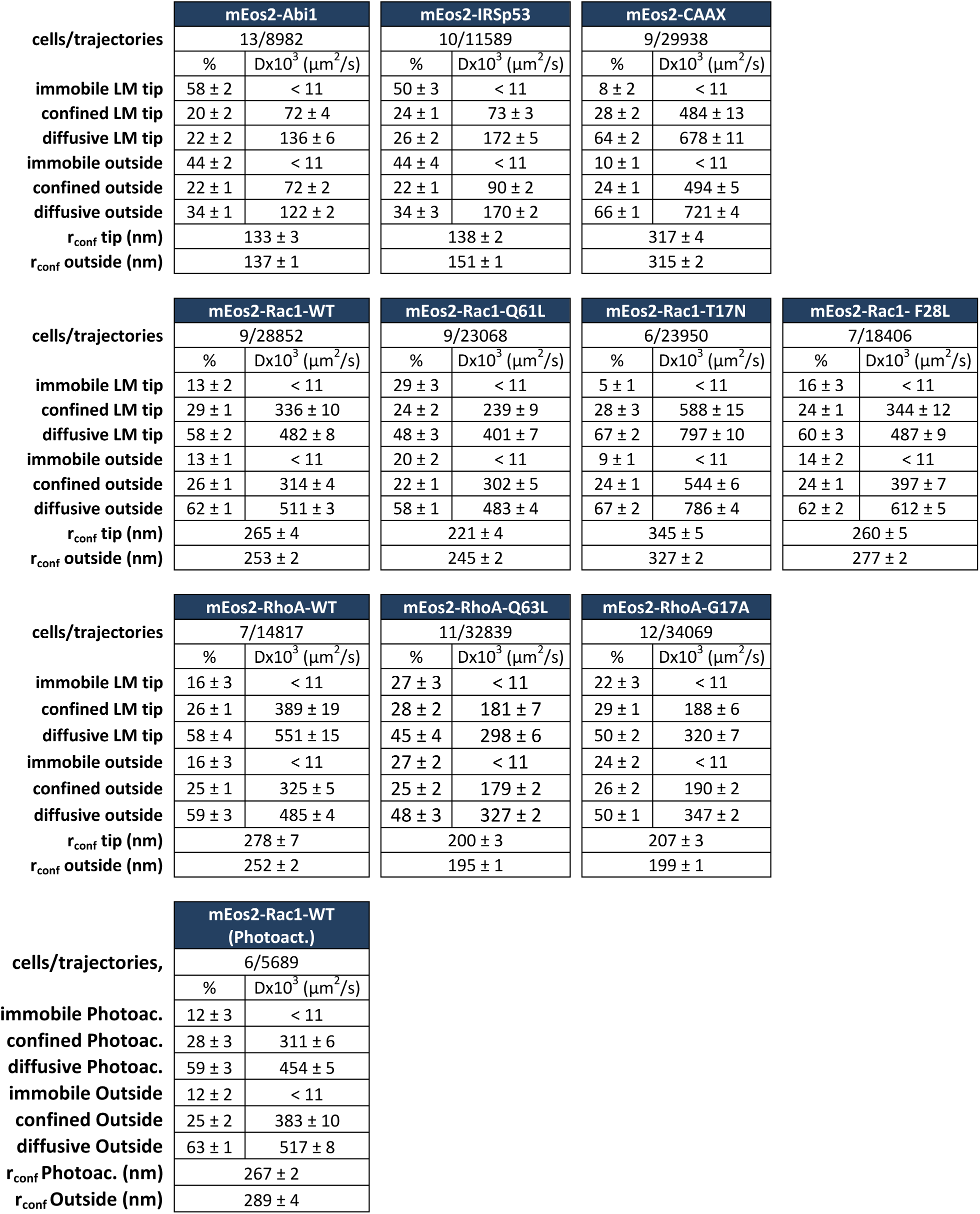
Results obtained using sptPALM acquisitions at 50 Hz (Figures 1, 2, 3 and 5) (LM, lamellipodium). Values correspond to Mean ± SEM.

**Table S2.** Results obtained using sptPALM acquisitions at 333 Hz (Figures 4) (LM, lamellipodium). Values correspond to Mean ± SEM.

|  | mEos2-Rac1-WT |  | mEos2-Rac1-Q61L |  | mEos2-Rac1-T17N |  | mEos2-Rac1- F28L |  |
| --- | --- | --- | --- | --- | --- | --- | --- | --- |
| cells/trajectories | 16/15648 |  | 14/15710 |  | 5/3767 |  | 11/4527 |  |
| | % | Dx10<sup>3</sup> ($\mu\text{m}^2/\text{s}$) | % | Dx10<sup>3</sup> ($\mu\text{m}^2/\text{s}$) | % | Dx10<sup>3</sup> ($\mu\text{m}^2/\text{s}$) | % | Dx10<sup>3</sup> ($\mu\text{m}^2/\text{s}$) |
| immobile LM tip | 16 ± 2 | < 52 | 40 ± 1 | < 52 | 12 ± 5 | < 52 | 24 ± 2 | < 52 |
| confined LM tip | 35 ± 1 | 623 ± 17 | 27 ± 1 | 363 ± 10 | 30 ± 3 | 812 ± 31 | 37 ± 2 | 448 ± 19 |
| diffusive LM tip | 50 ± 2 | 964 ± 19 | 33 ± 1 | 619 ± 13 | 58 ± 5 | 1 149 ± 26 | 39 ± 3 | 792 ± 26 |
| immobile outside | 9 ± 2 | < 52 | 21 ± 1 | < 52 | 13 ± 4 | < 52 | 18 ± 3 | < 52 |
| confined outside | 33 ± 1 | 695 ± 11 | 30 ± 1 | 443 ± 8 | 30 ± 3 | 760 ± 26 | 32 ± 2 | 502 ± 21 |
| diffusive outside | 58 ± 2 | 1 006 ± 11 | 49 ± 1 | 680 ± 9 | 57 ± 5 | 1 047 ± 23 | 50 ± 3 | 785 ± 23 |
| r<sub>conf</sub> tip (nm) | 162 ± 2 | | 124 ± 1 | | 180 ± 3 | | 139 ± 2 | |
| r<sub>conf</sub> outside (nm) | 165 ± 1 | | 134 ± 1 | | 176 ± 3 | | 145 ± 2 | |

|  | mEos2-CAAX |  |
| --- | --- | --- |
| cells/trajectories | 8/3824 |  |
| | % | Dx10<sup>3</sup> ($\mu\text{m}^2/\text{s}$) |
| immobile LM tip | 10 ± 1 | < 52 |
| confined LM tip | 28 ± 1 | 847 ± 48 |
| diffusive LM tip | 62 ± 1 | 1 102 ± 43 |
| immobile outside | 8 ± 1 | < 52 |
| confined outside | 36 ± 3 | 783 ± 26 |
| diffusive outside | 56 ± 4 | 1 131 ± 23 |
| r<sub>conf</sub> tip (nm) | 176 ± 4 | |
| r<sub>conf</sub> outside (nm) | 170 ± 2 | |

